# ROCCO: A Robust Method for Detection of Open Chromatin via Convex Optimization

**DOI:** 10.1101/2023.05.24.542132

**Authors:** Nolan H. Hamilton, Terrence S. Furey

## Abstract

**Motivation:** Analysis of open chromatin regions across multiple samples from two or more distinct conditions can determine altered gene regulatory patterns associated with biological phenotypes and complex traits. The ATAC-seq assay allows for tractable genome-wide open chromatin profiling of large numbers of samples. Stable, broadly applicable genomic annotations of open chromatin regions are not available. Thus, most studies first identify open regions using peak calling methods for each sample independently. These are then heuristically combined to obtain a consensus peak set. Reconciling sample-specific peak results *post hoc* from larger cohorts is particularly challenging, and informative spatial features specific to open chromatin signals are not leveraged effectively.

**Results:** We propose a novel method, `ROCCO`, that determines consensus open chromatin regions across multiple samples simultaneously. `ROCCO` employs robust summary statistics and solves a constrained optimization problem formulated to account for both enrichment and spatial dependence of open chromatin signal data. We show this formulation admits attractive theoretical and conceptual properties as well as superior empirical performance compared to current methodology.

**Availability and Implementation:** Source code, documentation, and usage demos for `ROCCO` are available on GitHub at: https://github.com/nolan-h-hamilton/ROCCO. `ROCCO` can also be installed as a standalone binary utility using [pip/PyPI](https://pypi.org/project/rocco/).

**Supplementary Information:** Supplementary material is available with this submission. Additional resources that may aid readers are available in the `ROCCO` GitHub repository.

## 1 Introduction

Nucleosomes, complexes of DNA and histone proteins, comprise the initial stage of chromatin compaction of the genome, reducing its occupying volume and enabling it to fit in cell nuclei [Li and Reinberg, 2011]. Most nucleosomal DNA is inaccessible for binding by transcription factors (TFs) that regulate gene expression. Genome-wide annotations of non-nucleosomal DNA, or open chromatin, therefore delineate where TFs can readily bind and effectively characterize the current gene regulatory program in a sample. Open chromatin landscapes vary across cell types and conditions, including in disease [Corces and Granja, 2018] reflecting cell and condition-specific gene regulation. To better understand this dynamic nature of gene regulation, the identification of open chromatin regions has become an important aspect of molecular studies of complex phenotypes.

Several assays have been developed for genome-wide measurement of open chromatin, including DNase-seq [Boyle et al., 2008] and ATAC-seq [Buenrostro et al., 2015]. These assays generate DNA fragments enriched for open chromatin regions that are then sequenced using short-read sequencers. Resulting reads are aligned to a reference genome, and regions with an enrichment of reads, or *peaks*, are identified as open chromatin. Peak calling is a necessary step as, unlike genes for which annotations are available for many species, there are not comprehensive, predefined standard databases of open chromatin regions.

Studies focused on determining changes in chromatin associated with differing cellular conditions or complex traits normally include many samples. For these studies, it is necessary to define a common set of open chromatin regions, or *consensus peaks*, to facilitate comparisons across sample groups. Typically, consensus peaks are determined by first annotating peaks independently in each sample. Then, these sample-specific peaks are merged based on one of several heuristics including: (1) Simply take the maximal set across all samples. This method, though, is particularly vulnerable to anomalous data since peaks from a single sample satisfy the inclusion criterion; (2) Include only peak regions that occur in *all* samples. This is usually too stringent due to variability in data quality across samples, especially when there is an expectation of differences; (3) Require that peaks be present in at least *M* = 1 … *K* samples, where the boundaries of the consensus peaks allow for some tolerance, *T*, for disparity in nucleotide position. Protocols in the spirit of this general method have been utilized in many open chromatin studies [Bao et al., 2015, Wang et al., 2018, Ming et al., 2020, Bentsen et al., 2020]. A difficulty in applying such methods is choosing appropriate *M* and *T* —a task manifesting rigid criteria that may ignore some open regions or may include spurious regions and that may not define well-supported peak boundaries.

More statistically sound methods have been developed for handling multiple samples. For the specific case of *K* = 2 samples, the Irreproducible Discovery Rate [Li et al., 2011] can be controlled to mitigate calling of irreproducible peaks. However, since most experimental designs include *K >>* 2 samples per group, it is difficult to apply this method broadly. Alternatively, Genrich^1^ offers a method for multiple samples in which *p*-values are combined using Fisher’s Method [Fisher, 1925]. While Genrich has been used successfully in several studies [Hofvander et al., 2019, Salavati et al., 2021, Tsaryk et al., 2022, Guerin et al., 2021], the independence assumption of Fisher’s Method may be problematic for large numbers of samples and/or in cases involving multiple technical replicates [Roy et al., 2019]. It also does not explicitly account for specific peak boundaries.

Here, we propose a novel method for identification of open chromatin regions across multiple samples, ROCCO: “**R**obust **O**pen **C**hromatin detection via **C**onvex **O**ptimization.” This method offers several favorable features:

- Accounts for both enrichment *and* spatial characteristics of open chromatin signals, the latter of which is an informative but often ignored aspect of ATAC-seq data that can be used to not only better detect regions but also improves on annotating peak boundaries;
- Leverages data from multiple samples without imposing arbitrary “hard” thresholds on a minimum number of samples declaring peaks;
- Is efficient for large numbers of samples;
- Does not require training data or a heuristically determined set of initial candidate regions, which are hard to define given the lack of a priori sets of validated open chromatin regions;
- Employs a mathematically tractable model granting useful performance guarantees.

We formally describe the algorithm utilized by `ROCCO` for the consensus peak problem and present a theoretical analysis. We then conduct several experiments to investigate `ROCCO`’s efficacy empirically, using a set of 56 samples from human lymphoblastoid cell lines.

**Table 1:** Notation Reference.

| Symbol | Description | Default Value |
| --- | --- | --- |
| $b$ | budget threshold on selected loci | .035 |
| $c_1, c_2, c_3$ | weights for score function $\mathcal{S}(i)$ | 1.0 |
| $f$ | objective function | N/A |
| $\gamma$ | penalty weight in (3) | 1.0 |
| $K$ | number of input samples | N/A |
| $\ell_i$ | decision variable for $i^{\text{th}}$ locus | N/A |
| $\boldsymbol{\ell}$ | vector of decision variables | N/A |
| $L$ | fixed interval size of input signals | 50 |
| $\mathcal{L}$ | contiguous genomic region | N/A |
| $n$ | number of loci | N/A |
| $N$ | max. randomized rounding iterations | 50 |
| $\mathcal{S}(\cdot)$ | locus score function | N/A |
| $\mathbf{S}_{\mathcal{L}}$ | $K \times n$ signal matrix over $\mathcal{L}$ | N/A |
| $\tau$ | median enrichment threshold in $\mathcal{S}(\cdot)$ | 0 |

## 2 Algorithm

We begin by introducing notation used throughout this manuscript and describing the structure of signal data used by `ROCCO` to detect accessible chromatin.

### 2.1 Notation and Definitions

Let *ℒ* be a contiguous genomic sequence, e.g. a chromosome, divided into *n* fixed-width loci, each consisting of *L* nucleotides as in Figure 1. For each sample *j*, we assume access to a signal, *s*_*i*_^*j*^, computed as a function of observed enrichment based on sequence reads at the *i*^th^ locus. For *K* total samples, this yields a *K × n* signal matrix used as input to `ROCCO`:

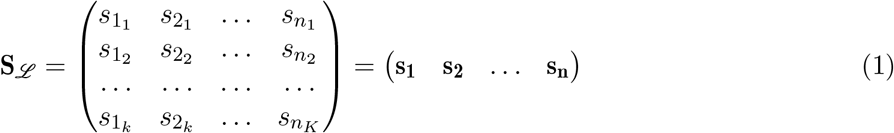

with **s**_**i**_ ∈ ℝ^*K*^ denoting the column vector of signal values among samples at the *i*^th^ locus. This matrix can be generated with a variety of methods, but a context-specific tool, `rocco prep` is included as a subcommand in the software implementation for convenience: Given a directory of samples’ BAM files, **S**_*ℒ*_ is generated with multiple calls to PEPATAC’s [Smith et al., 2021] `bamSitesToWig.py` tool.

**Figure 1:**
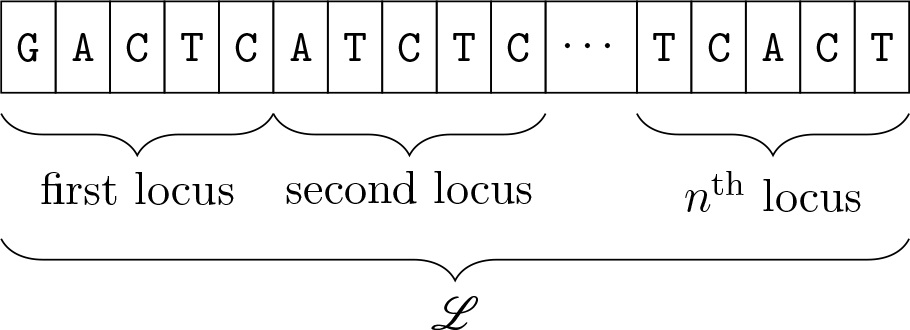
Genomic region *ℒ* consisting of *n* fixed-width loci containing *L* = 5 nucleotides each.

### 2.2 Scoring Loci

To determine consensus open chromatin regions, we first score each locus while accounting for enrichment (*g*_1_), dispersion among samples (*g*_2_), and a measure of local volatility in enrichment (*g*_3_).

Specifically, we take *g*_1_(*i*) to be the median, and *g*_2_(*i*) to be the median absolute deviation [Pham-Gia and Hung, 2001] of the *K* signal values at the *i*^th^ locus:

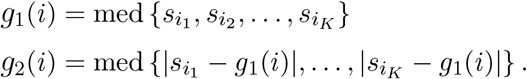

Large *g*_1_ and low *g*_2_ correspond to regions of high enrichment with little dispersion among samples— a favorable combination of traits to emphasize when predicting accessibility. We also leverage the disparities between enrichment signal values at adjacent loci, normalized by the current locus’s enrichment,

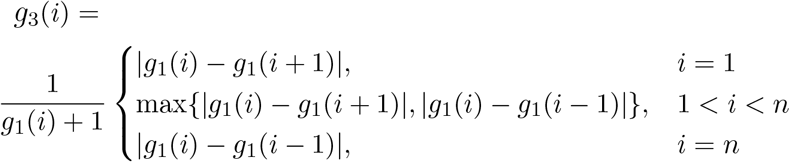

A fundamental aim of *g*_3_ is to more precisely annotate the edges and immediately adjacent regions of peaks where signals may be low before or after an abrupt shift in enrichment characterizing the nearby peak. See Figure 2 for a visual demonstration on an idealized, continuous enrichment signal.

**Figure 2:**
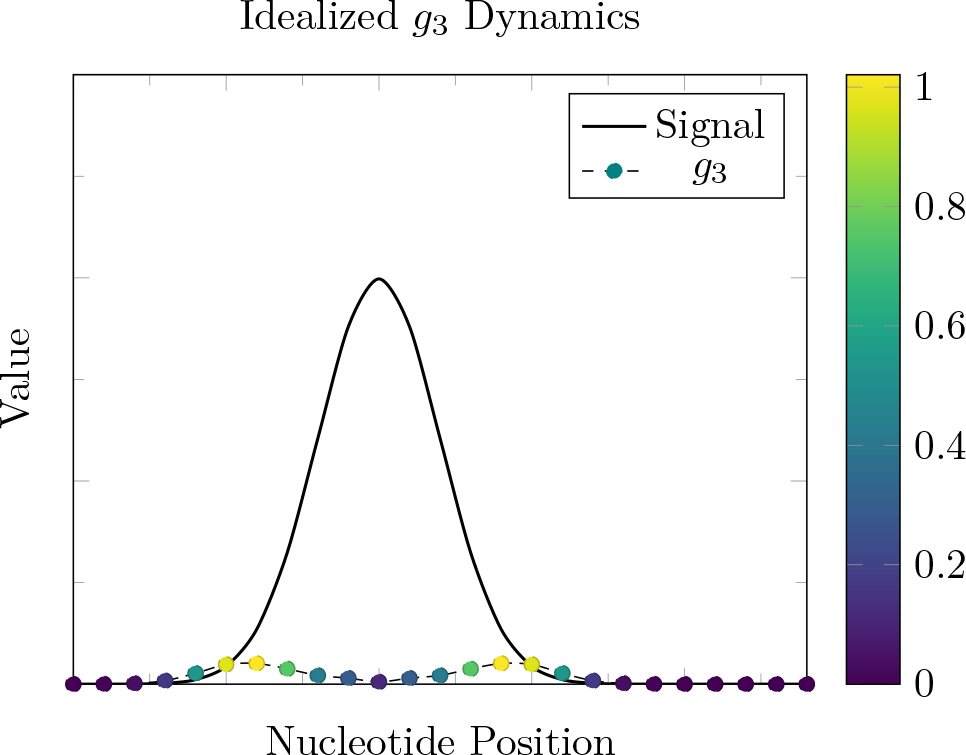
*g*_3_ is designed to more accurately determine peak edges to capture enriched regions in their whole. This function is greatest (yellow marks) near the ends of the enriched region and lowest (dark blue marks) at peak centers in this idealized example.

In (2), we define the score piece-wise and take a simple linear combination of *g*_1_, *g*_2_, *g*_3_—or set this score to 0 if median enrichment is below a defined threshold. This enrichment threshold can encourage sparsity that may have favorable implications for computational efficiency during optimization and mitigate consideration of regions that are unlikely accessible.

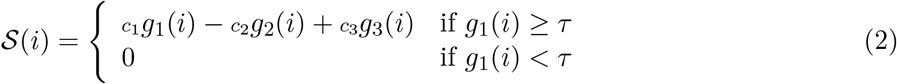

where *τ≥*0 is the minimum enrichment threshold placed on the median signal, and *c*_1_, *c*_2_, *c*_3_*≥*0 are coefficients for each term. By default, each scoring term has the following weights: *c*_1_, *c*_2_, *c*_3_ = 1, and *τ* = 0. These defaults provide strong performance but can be modified in the software implementation of ROCCO to suit users’ specific needs. For example, users desiring more conservative peak predictions may wish to set *τ >* 0.

𝒮 (*i*) *≤*0 does not necessarily preclude the corresponding locus from being selected within a consensus peak, since the objective function (See Eqn. 3) is not completely dependent on the locus score. Note that for default parameters, *𝒮* (*i*) *≥* 0.

### 2.3 Optimization

We address open chromatin detection as a constrained optimization problem. Let ***ℓ*** *∈{*0, 1*}*^*n*^ be a vector of binary decision variables, where we label *ℓ*_*i*_ = 1 if the *i*^th^ locus is present in open chromatin, and *ℓ*_*i*_ = 0 otherwise.

We impose a budget constraint *upper-bounding* the proportion of selected loci in a given input chromosome. Let *b ∈* [0, 1] be the maximum percent of loci that can be selected, i.e.,

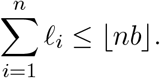

This constraint controls sensitivity in peak predictions and prevents unrealistic solutions in which an excessive fraction of chromatin is declared open. With estimates for the fraction of accessible chromatin hovering around 3*−*4% of the human genome [Song et al., 2011, Sahinyan et al., 2022], we accordingly set *b* = .035 as the default value. Since the budget applies to each chromosome independently, and the accessibility for each chromosome can vary, the software implementation allows for chromosome-specific parameters to be defined.

To optimize selection of accessible regions, the following objective function is *minimized*:

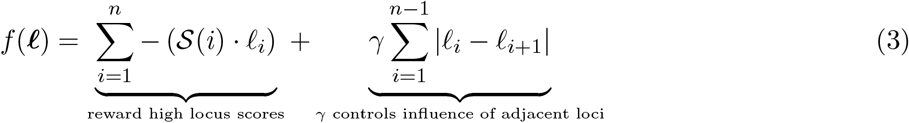

The first term rewards loci with high 𝒮 scores, e.g. those with consistently high enrichment across samples or those on the edges of greatly enriched regions. The second term is introduced to account for spatial proximity of loci during optimization and controls the influence of signals in adjacent loci: For a given budget *b*, as *γ* is increased corresponding to a greater influence of adjacent regions, fewer but longer *distinct* regions are annotated as open, yielding simpler solutions in a topological sense. This pattern is exhibited in Figure 3. To incorporate the described objective and budget constraint, we pose the following constrained optimization problem:

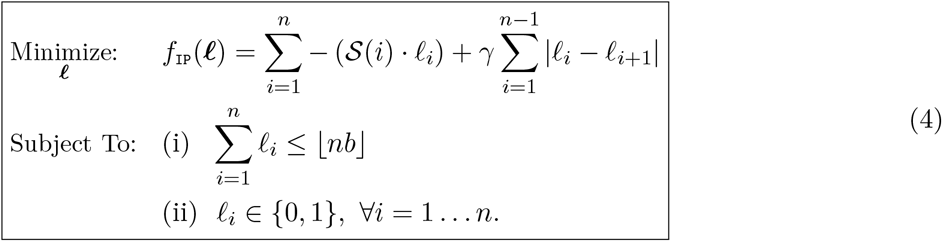

**Figure 3:**
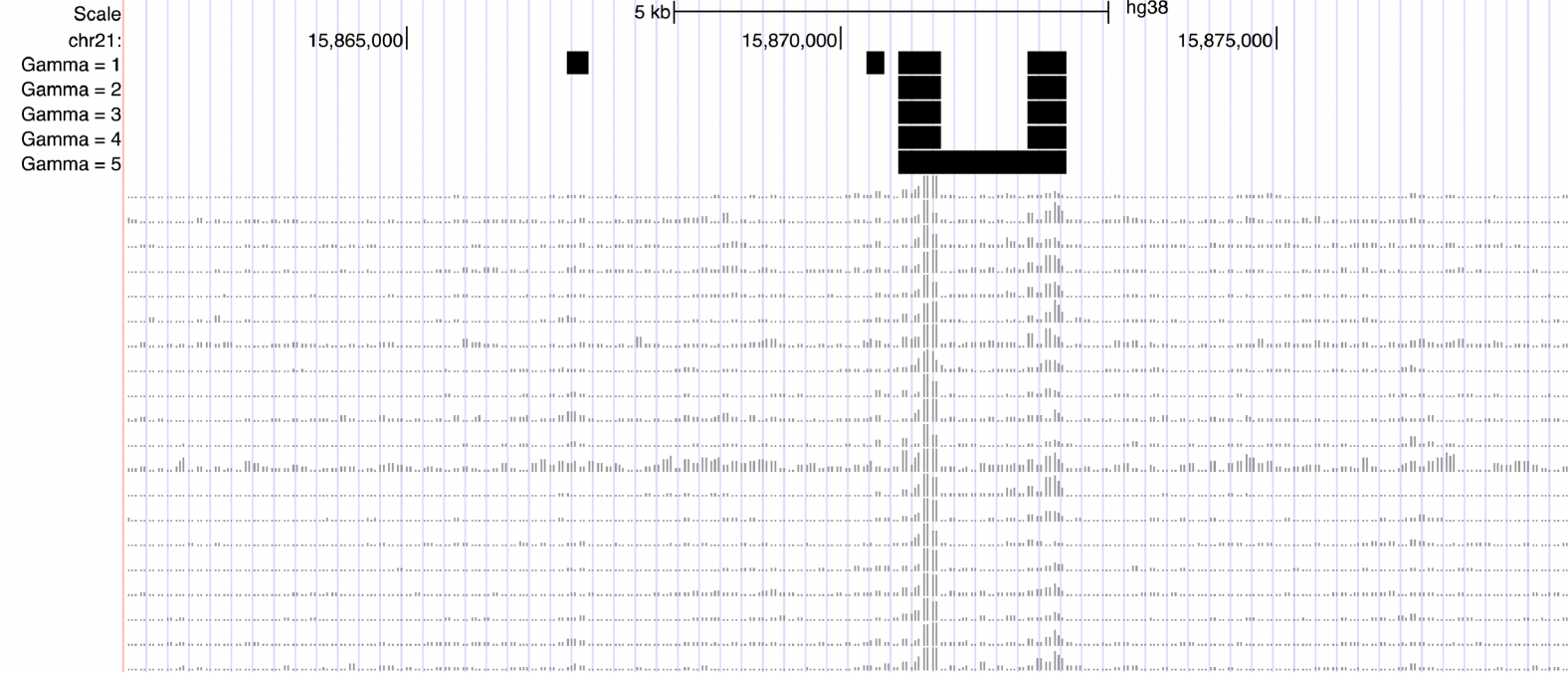
Example behavior of `ROCCO` in the UCSC Genome Browser [Karolchik et al., 2003] as *γ* is increased. The black bars in each track correspond to `ROCCO`’s predictions given the replicate signal data below. In the last row, we see that two distinct regions of enrichment are merged due to the strong influence of adjacent loci imposed by the *γ* = 5 parameter.

Constraint (ii) restricts the feasible region to integer solutions. In general, such constraints yield difficult optimization problems, for example, because gradients are not defined for functions over the integers and convexity cannot be leveraged. Indeed, general integer programming is known to be NP-hard [Korte and Vygen, 2012]. A common remedy is to convert the original, integer-constrained formulation to an analogous problem with convenient analytic properties. Accordingly, we substitute the constraints

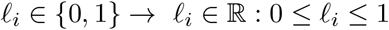

to obtain the following convex optimization problem:

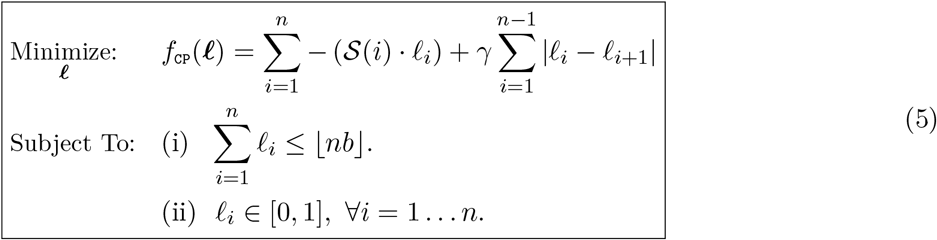

As we will see, this formulation maintains the essence of the original problem in (4) and confers several useful properties. In general, convexity is a highly-valued feature in constrained optimization as it ensures every local minimum is also a global minimum, thereby preventing instances of “premature” convergence to suboptimal solutions [Boyd and Vandenberghe, 2004].

#### Theorem 1.

*The problem in (5) can be solved in polynomial time for a globally optimal solution*.

Linear programs (LPs) are a special class of convex optimization problems in which both the objective and constraints are linear functions of the decision variables. Though general convex problems can often be solved efficiently in practice, there are certain intractable instances. In contrast, LPs can be solved in worst-case polynomial time with respect to the number of variables [Boyd and Vandenberghe, 2004]. Accordingly, the proof of Theorem 1, deferred to Supplementary Material 1.1, relies on showing that an optimal solution to (5), ***ℓ*** ^*CP*^*∈*ℝ^*n*^, is obtained from the *n*-dimensional truncation of the optimal solution to the following LP:

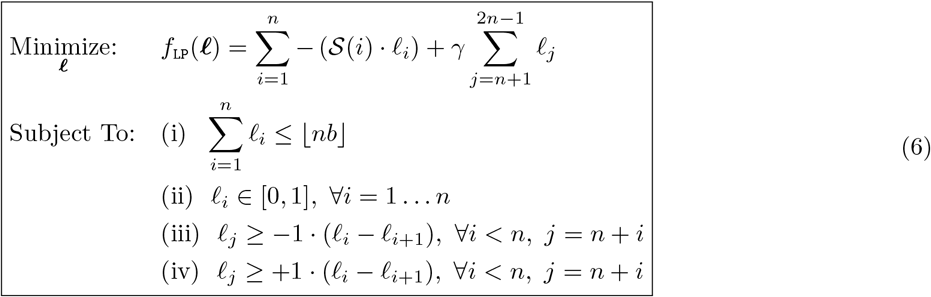

which we denote as ***ℓ***^*LP*^ *∈* ℝ^2*n−*1^. Moreover, the optimal objective values for (5) and (6) are equal:

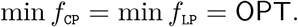

The time complexity of standard interior-point methods for solving LPs with *n* decision variables and a *d*-bit data representation is 𝒪 (*n*^3^*d)*, though several methods with improved worst-case bounds have been proposed [Karmarkar, 1984, Vaidya, 1989, den Hertog, 1994]. It is important to note that, in practice, modern solvers offer much greater efficiency than this worst-case bound might suggest by exploiting problem structure [Koch et al., 2022, Boyd and Vandenberghe, 2004]. Indeed, (6) possesses a particularly sparse objective and constraint matrix that allows for surprising efficiency showcased in Supplementary Material 1.3.

After solving the relaxed form of an integer program, it is often necessary to refine the solution for feasibility in the original integer-constrained region. Exact combinatorial techniques, such as branch and bound, can be applied to find optimal integer solutions; However, such methods may incur prohibitive computational expense and do not offer polynomial runtime guarantees. It is often more practical to use an approximation scheme [Williamson and Shmoys, 2011].

We find in our own experiments that solutions to (6) are nearly integral (Figure 4) and can be rounded immediately, for instance, with the floor function, to obtain a feasible solution without a substantial sacrifice in practical performance. However, this near-integrality cannot be guaranteed in general, and a more robust procedure is preferred, which we now describe.

**Figure 4:**
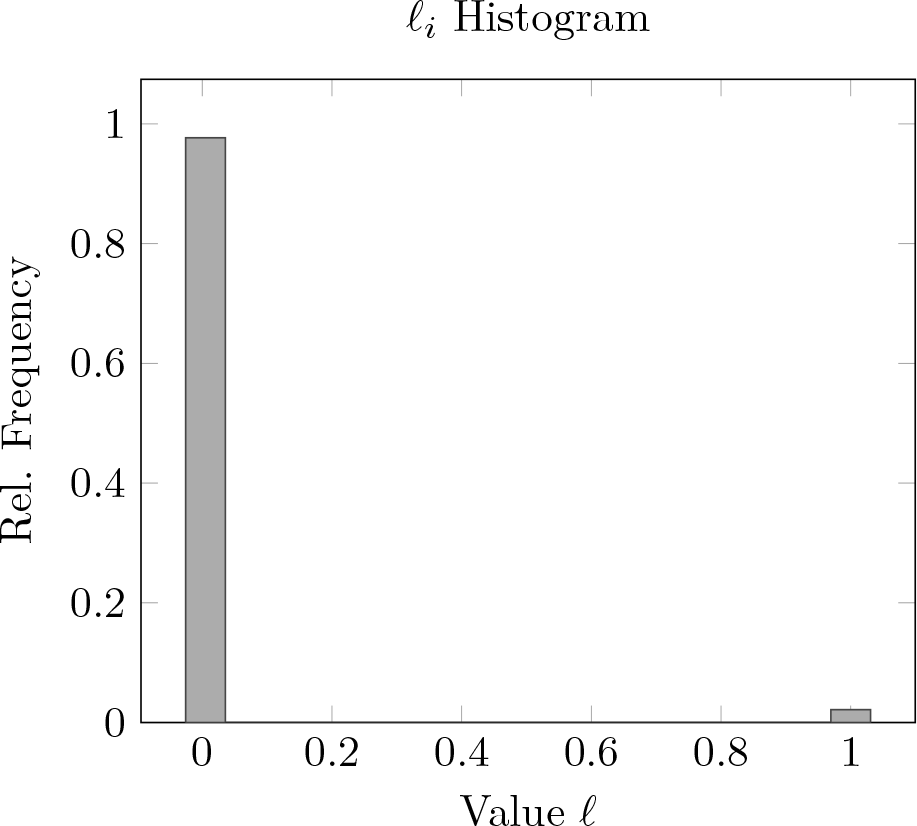
Observed distribution of decision variables after solving (6) with budgets *b ∈{*.01, .025, .05, .075, .10*}* on 50 random subsamples (*K* = 40) of the ATAC-seq data detailed in Section 5.2 and pooling the solutions from each run. See attached material for exact relative frequencies used to generate this plot.

#### Algorithm 1: Drawing *ℓ* ^*rand*^

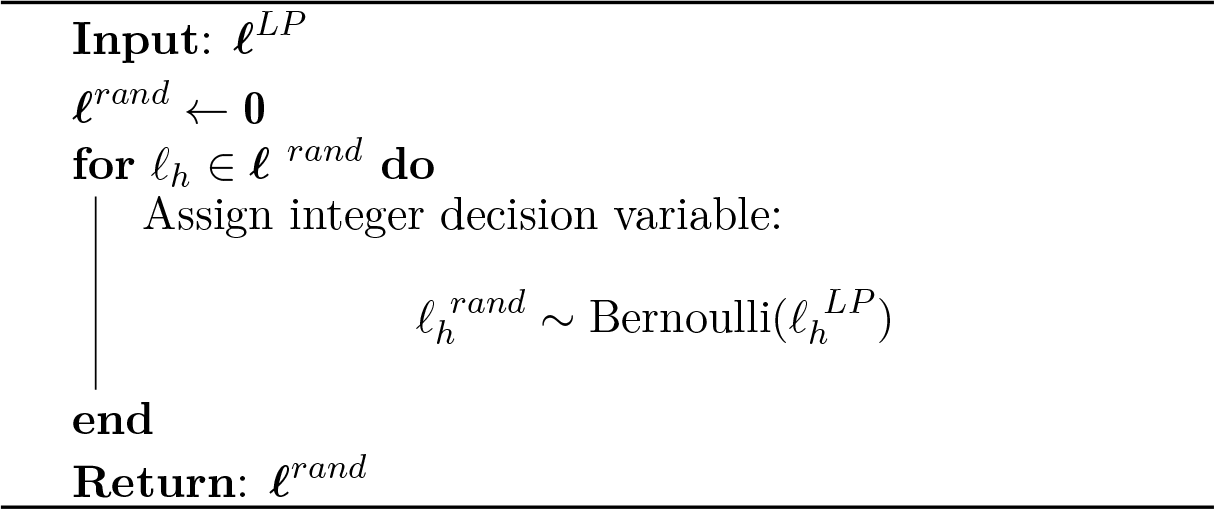

Given a solution to the relaxed formulation (6), we devise a procedure, denoted `RR`, based on randomized rounding [Raghavan and Tompson, 1987], to obtain a set of candidate integral solutions, L_*N*_ . This set is generated by executing *N* iterations of Algorithm 1, after which `RR` picks the best feasible solution with the lowest objective value.

Note that in a given solution space *ℱ∈*ℝ^*D*^, *D ≥*1, integral solutions cannot yield better performance than the best real-valued solution. The set of integral solutions is a proper subset of *ℱ* . For this reason, the quality of integer solutions can be judged with reference to *f*_LP_(***ℓ***^*LP*^). In light of this, the construction of solutions ***ℓ*** ^*rand*^*∈***L**_*N*_, with each 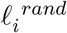 defined as a Bernoulli-distributed random variable with parameter 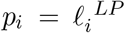, grants convenient properties arising from linearity of expectation. Namely,

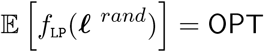

with constraints satisfied in expectation by ***ℓ*** ^*rand*^. We can use these expected values and leverage concentration inequalities to make probabilistic assertions regarding the solutions present in **L**_*N*_.

#### Theorem 2.

*Let* L_*N*_ *be a set of N ≥* 1 *random solutions generated with Algorithm 1, and let c >* 1, *a >* 0 *be real numbers satisfying* 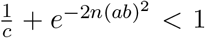 *for n loci and budget b ∈* (0, 1). *Then with high probability*, **L**_*N*_ *contains at least one solution with both (a) an objective value no more than c ·* `OPT` *and (b) no more than nb*(1 + *a*) *loci selected*.

In short, Theorem 2 is proven^2^ using Markov’s and Hoeffding’s inequalities to show that the probability of one or more ***ℓ***^*rand*^ *∈* **L**_*N*_ satisfying both criteria is at least

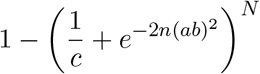

which quickly approaches 1 for increasing *N* . We emphasize that this expression is a lower bound on the probability of a satisfying solution, and we often observed multiple such solutions in **L**_*N*_ during the course of our experiments. But Theorem 2 allows us to make more general assertions under the supposition 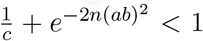. Since *n* is on the order of millions for default locus size *L* = 50, this criterion is satisfied even for quite small *c >* 1, *a >* 0.

The `RR` procedure has linear time and space complexity, making it a minor contributor to overall computational expense. Though this random procedure technically renders `ROCCO` stochastic, for the default *N* = 50 `RR` iterations, we observed only minor variation in solutions returned from independent runs of `ROCCO`. As seen in Supplementary Material (Table S3), a pairwise Jaccard similarity matrix for five independently-generated `ROCCO` solutions contains values no less than 0.9977.

Algorithm 2 offers a pseudocode representation of `ROCCO` as a whole. The object returned by Algorithm 2 is an *n*-dimensional decision vector, ***ℓ***^*∗*^, used to select loci as accessible. Note, contiguous selections (that is, sequences of loci such that 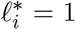) are merged into single peaks in the final BED file.

#### Algorithm 2: ROCCO

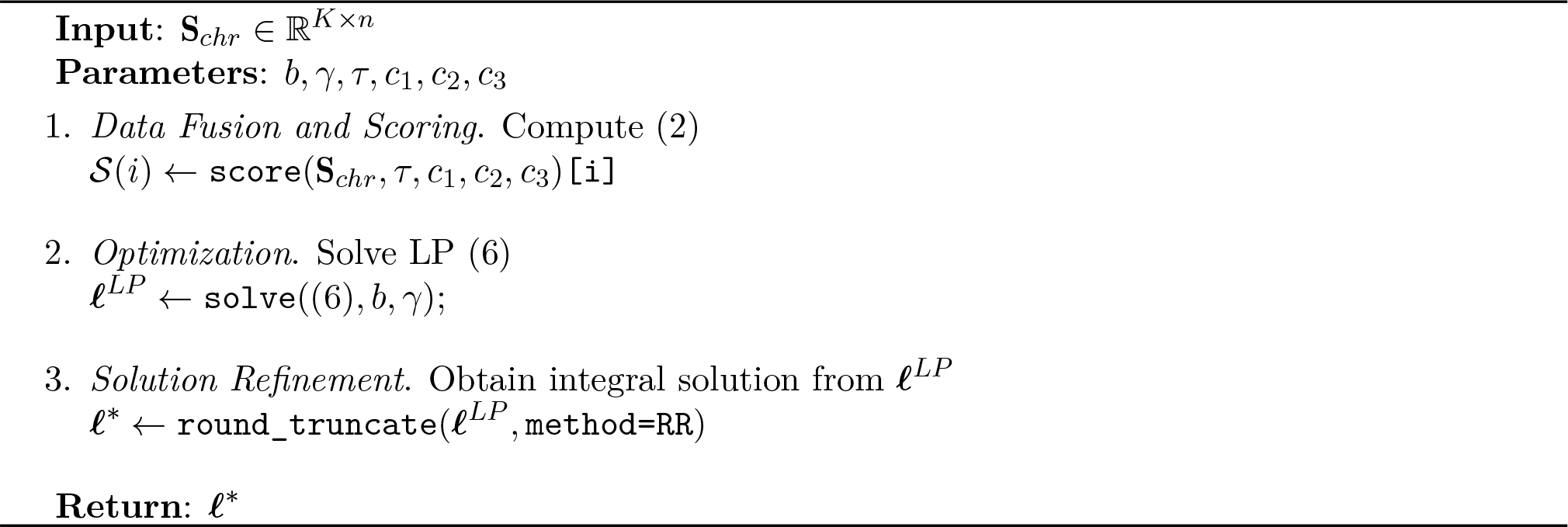

#### Remark 1.

*Large Sample Sizes*. Note that only the scoring step is directly affected in runtime by the number of input samples *K*, running on the order of *nK* elementary operations using the median of medians algorithm. Practical scenarios satisfy *K << n*, making the number of input samples a minor contributor to computational expense during the calculation of 𝒮 (·). Perhaps more importantly, because the scoring step is asymptotically dominated in worst-case computational expense by the optimization step, which is independent of *K*, the worst-case time complexity of `ROCCO` is likewise independent of the sample size, *K*.

## 3 Results

We performed several experiments to assess `ROCCO`’s detection performance using ATAC-seq data from 56 human lymphoblast samples (see Section 5). Additional analyses and details are available in the Supplementary Material.

### 3.1 Detection Performance

A noteworthy limitation in experiments comparing performance of open chromatin detection methods is a lack of high-confidence annotations against which to test. However, to gauge performance and ensure viability, some proxy for ground truth is needed. Following [Zhao and Boyle, 2021], we constructed a *union set*, `GT`, of conservative Irreproducible Discovery Rate (IDR) [Li et al., 2011] peaks from ENCODE *transcription factor* ChIP-seq experiments in the `GM12878` lymphoblast cell line. We assume that the majority of annotated transcription factor binding sites will correspond to open chromatin regions, but we note that variability in binding at a snapshot in time, the incomplete annotation of all transcription factor binding, and cases where factors can bind to non-accessible chromatin introduce notable limitations. But we argue that this data is sufficient to compare the relative performance of distinct methods. The ℱ_*β*_-score, defined below, was then used to assess the ability to recover and bound regions in `GT` using ATAC-seq data from the 56 independent samples. Details regarding the construction of the `GT` data set can be found in Supplementary Material 1.7.

For each method, we generated consensus peaks using previously determined alignments for the *K* = 56 samples. We then computed precision as

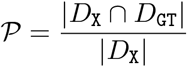

and recall as

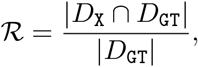

where *D*_*X*_ denotes the consensus peaks obtained from method *X* and set intersections in the numerators are computed using `bedtools intersect`. The *F*_*β*_-score was then calculated as the harmonic mean of precision and recall where recall is weighed *β* times as much as precision, that is:

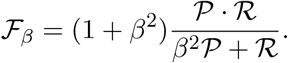

As in [Zhao and Boyle, 2021], we use the ℱ_*β*_-score as the primary metric for comparison of methods since it intuitively combines both precision and recall and is less affected by extreme regions of the precision-recall curve that do not correspond to realistic use-cases.

#### 3.1.1 Detection Performance: Benchmark Methods

Most methods to determine consensus peaks begin by identifying sample-specific peaks. For this step, we employed the widely-used MACS2 software [Gaspar, 2018] using parameters commonly specified for ATAC-seq experiments (see Section 5). With these, we used a common heuristic to specify consensus peaks [Yang et al., 2014]. Namely, `MACS2-Consensus` only retained merged peaks supported by a majority of samples with a 100bp tolerance in chromosome position across samples. Genrich is another method for consensus peak calling we tested that analyzes samples separately, calculating *p*-values for each. It then applies Fisher’s method to combine *p*-values at each genomic region. We also generated peak sets with `MACS2-Pooled`, which combined alignments from all samples into one BAM file and then used MACS2 to call peaks on this combined alignment file.

See Supplementary Material 1.8 for exact configurations used to produce results for these MACS2-based methods and for Genrich.

#### 3.1.2 Detection Performance: Results

For an initial visual comparison of methods, Figure 5 displays peak calls from each in 100kb and 20kb regions on chromosome 19 in the UCSC Genome Browser [Karolchik et al., 2003]. We also include ATAC-seq signals from 25 of the lymphoblast samples being evaluated. As expected, all methods identify regions with consistently strong signals across all samples. They vary, though, in the contiguity and boundaries of these regions. There are also method-specific regions, as well as ones called by multiple, but not all, methods.

**Figure 5:**
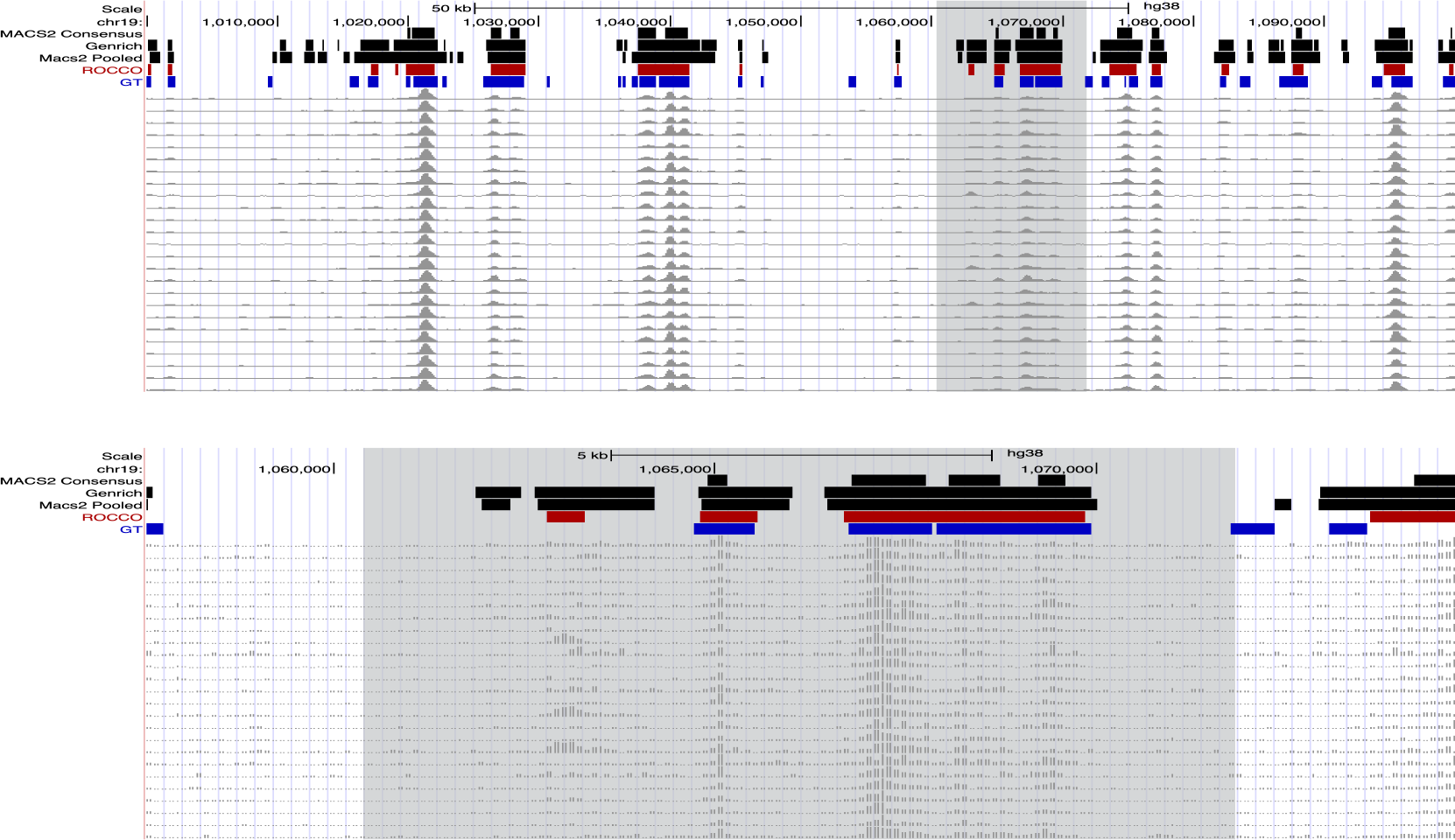
*Example Behavior over* chr19:1000000*−*1100000. Consensus peak calls from each method tuned using the ℱ_*β*_ score (*β* = 1.0), displayed at resolutions of 100,000 nucleotides (top) and 20,000 nucleotides (bottom)

To quantify genome-wide detection performance of the methods, we evaluated across several values

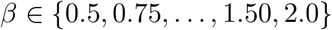

to address a plausible but encompassing range of recall/precision prioritizations. The most extreme cases *β* = 0.5, *β* = 2.0 were included for completeness but may not be particularly well-motivated by realistic usage since the corresponding ℱ_*β*_ score can be unduly improved by simply rejecting any uncertain predictions or accepting all plausible predictions, respectively.

For each *F*_*β*_-score, we tuned each method over a range of significance thresholds deemed reasonable given their underlying models to maximize their performance. For `Genrich`, we tested

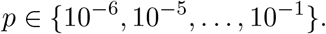

For the MACS2-based methods, we tested:

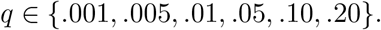

For `ROCCO`, the budget parameter is most fundamental and upper-bounds the fraction of genomic region *ℒ*that can be selected. We thus use *b* as the tuning parameter for `ROCCO`, leaving *γ, τ, c*_1,2,3_ as their default values, and evaluated:

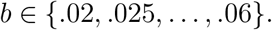

For each *β* listed above, given a method *X* and parameter *r*, we computed tuned performance as

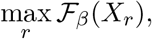

that is, the best-observed performance of the method while sweeping its most fundamental parameter. These values are recorded in Table 2. `ROCCO` matched or exceeded the best performance of every benchmark method for all six *β* values. The performance disparity between `ROCCO` and the second best-scoring method was smallest for the most recall-dependent case *β* = 2, which we have stated is only partially informative and particularly vulnerable to spurious predictions.

**Table 2:** Performance for each method is recorded after tuning for the *F*_*β*_ value in the leftmost column.

| $\beta$ | ROCCO | Genrich | MACS2-Pooled | MACS2-Consensus |
| --- | --- | --- | --- | --- |
| 0.50 | **0.651** | 0.460 | 0.390 | 0.643 |
| 0.75 | **0.596** | 0.489 | 0.427 | 0.586 |
| 1.0 | **0.579** | 0.520 | 0.465 | 0.545 |
| 1.25 | **0.580** | 0.545 | 0.501 | 0.517 |
| 1.50 | **0.582** | 0.566 | 0.531 | 0.498 |
| 2.0 | **0.603** | 0.595 | 0.577 | 0.475 |

As mentioned above, `ROCCO` allows for specifying chromosome-specific parameters to account for differences in accessibility across the genome. For a cursory investigation into the effects of this practice, we tuned the budget parameter for ℱ_1_ via grid search for each chromosome. We observed a non-trivial increase in performance (*F*_1_ = .620) compared to a constant budget for all chromosomes (*F*_1_ = .579 as in Table 2), but we expect additional improvements from a more rigorous, technically sound approach in which budgets are not restricted to an arbitrary set of values.

A comparison of methods using their *default* significance thresholds without tuning is included in the Supplementary Material S2, where `ROCCO` offers the greatest ℱ_*β*_-score in all but the *β* = 0.5, *β* = 2.0 experiments. Likewise, Supplementary Material 1.3 includes experiments assessing `ROCCO`’s computational efficiency in both theory and practice.

### 3.2 Variation in Sample Size/Quality

Ideally, a consensus peak calling procedure will

- Effectively leverage data presented by multiple samples.
- Yield robust results in the presence of varying sample quality and size.
- Scale efficiently for large sample sizes.

In this section, we conduct several analyses to consider these aspects for `ROCCO`.

In the first experiment, `ROCCO` is repeatedly executed using random subsamples of *K*_sub_ ∈ {5, 10, 15, …, 50} ATAC-seq alignments from the data set in Section 5.2 as input. The subsamples’ respective output peak sets are then compared to the peak set obtained by running `ROCCO` on the full set of *K* = 56 ATAC-seq alignments. To compute similarity between the subsamples’ peak sets and the entire sample’s, we measure the Jaccard index between their respective BED files using `bedtools` [Quinlan and Hall, 2010]. The average Jaccard indexes for each *K*_sub_ are recorded in Figure 6 along with 95% confidence intervals. Notably, with only *K*_sub_ = 5 samples, `ROCCO` generated peak sets roughly 70%-similar to the `ROCCO`’s peak set generated using all *K* = 56 samples. Moreover, `ROCCO` produced strictly increasing Jaccard indexes for increasing subsample sizes, indicating an effective utilization of additional samples. Though the purpose of this experiment is to evaluate `ROCCO`’s approximation of the full-sample-derived results with respect to varying subsample sizes, we note that detection performance as measured in Section 3.1 likewise improved with respect to increasing *K*_sub_ from ℱ_1_ = 0.546*±*0.011 to ℱ_1_ = 0.5783*±*.001.

**Figure 6:**
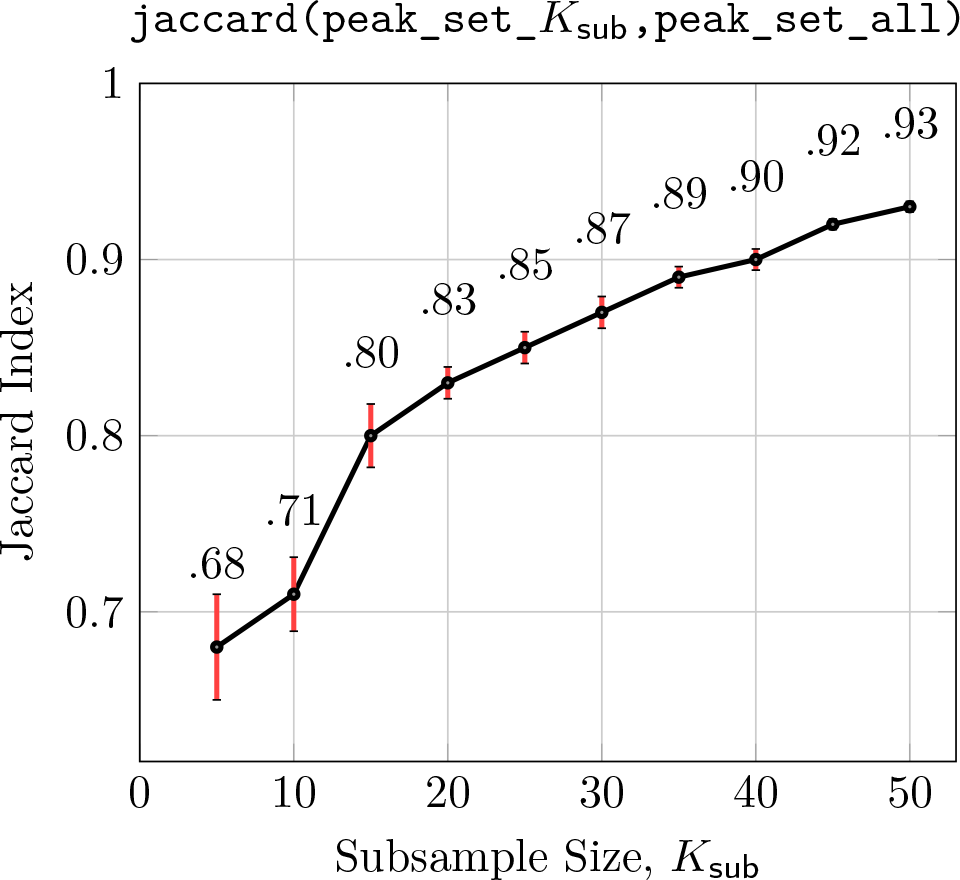
In fifty experiments for each *K*_sub_*∈{*5, 10, 15, …, 50*}, K*_sub_ ATAC-seq alignments are randomly subsampled and supplied as input to `ROCCO`. The fifty resulting output BED files are used to compute the average Jaccard index (95% C.I.) to `ROCCO`’s results obtained using all *K* = 56 samples.

Regarding efficiency, we note that the cpu-time required to execute `ROCCO` genome-wide was affected negligibly by *K*_sub_, with the average runtime for *K*_sub_ = 5 and *K*_sub_ = 50 differing by less than 20% despite the 1000% increase in samples. This result is informed theoretically by Remark 1, where the time complexity of `ROCCO` is shown to be asymptotically independent of sample size *K*.

The second experiment compares the effect of data quality on consensus peak sets generated by executing `ROCCO` independently on the ten best and worst samples as measured by the transcription start site (TSS) enrichment score [Smith et al., 2021]. The data from the 56 lymphoblastoid samples are of relatively good quality (Supplementary Material 1.6), as evidenced by minimum TSS enrichment score of 4.95. Nonetheless, the distribution of scores reflects appreciable differences in sample quality between the left and right tails. With this considered, the relatively small disparities in `ROCCO`’s detection performance shown in Figure 7 indicate robustness to variation in sample quality. In comparison, `MACS2-Consensus`, the best-performing alternative method in this experimental setting, returns lower ℱ_1_-scores for both the worst ten samples (ℱ_1_ = 0.492) and best ten samples (ℱ_1_ = 0.530) and a larger disparity between performance in each case.

**Figure 7:**
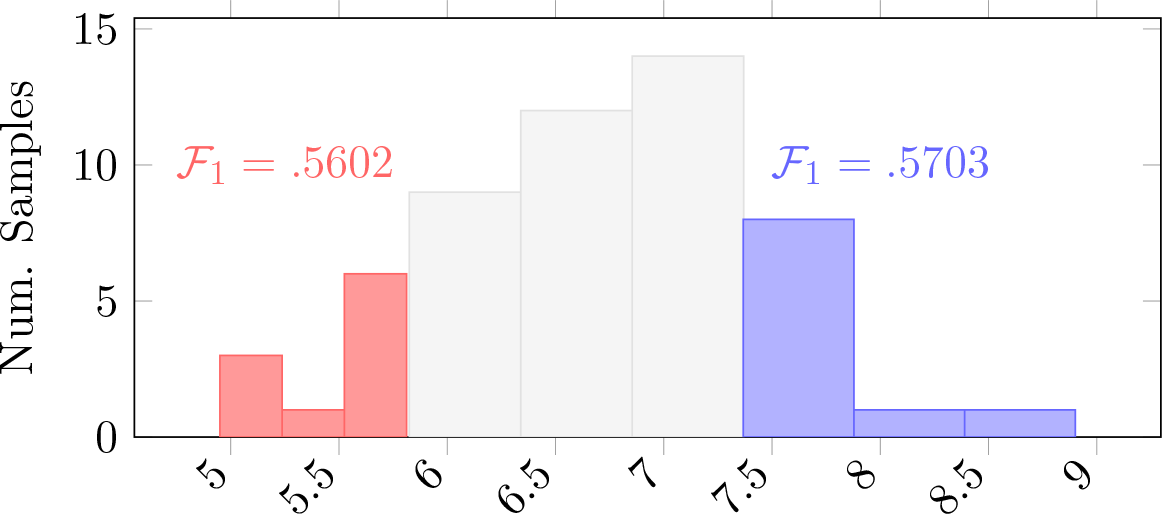
Histogram of TSS enrichment scores [as defined by ENCODE](https://www.encodeproject.org/data-standards/terms/) for the *K* = 56 ENCODE ATAC-seq samples used in experiments. `ROCCO` was run twice with default parameters—once using the *K* = 10 *worst* samples (red) and again using the *K* = 10 *best* samples (blue) as measured by TSS score.

### 3.3 Differential Accessibility Testing with ROCCO

A key motivation for the development of `ROCCO` was for experimental designs where ATAC-seq data is generated from multiple samples within two or more distinct groups. Using these data, a key question regards the location of genomic regions over which accessibility differs significantly between groups. Knowledge of such regions may yield insights into regulatory mechanisms responsible for phenotypic differences. To offer a template for differential analysis with `ROCCO`, we ran a simple experiment comparing the accessibility landscapes of males and females in the lymphoblast data described in Section 5.2.

To ensure that peaks specific to a particular group are considered, `ROCCO` was run independently on each group’s respective samples and then merged to produce the final set of consensus peaks. The software implementation of `ROCCO` offers a convenient means to separate samples by group for peak calling, the merging of peaks, and the generation of a count matrix across all samples that can be used for input to differential detection methods such as DESeq2 [Love et al., 2014]. Note, a Jupyter Notebook tutorial addresses steps for differential analysis with `ROCCO` and is available on the GitHub repository.

In this demonstration experiment, `ROCCO` was run independently on the 23 female and 33 male lymphoblastoid cell lines using default parameters, and the peaks were merged to create a final set of*∼*172,933 consensus peaks. We note that this merged set included 23,865 peaks only detected in the male samples and 19,165 peaks only detected in the female samples. Further, there were ~17,000 peaks in each of these sets that were not included in the consensus set derived from all samples combined. This underscores the importance of calling group-specific peaks in the context of differential analysis between imbalanced samples.

Despite the seemingly large number of phenotype-specific peaks, our previous experience with these analyses has shown that most of these are due to low signal peaks that are not significantly different across groups. Therefore, DESeq2 was then used to detect significant differentially accessible regions across the merged consensus peaks. At FDR-adjusted *p <* 0.05, 3,141 peaks spanning 2,275,100 base pairs were identified as differentially accessible. About 93% (2,916) of these peaks were observed in chromosome X, which is unsurprising given the division of samples into male and female groups.

## 4 Discussion

In this manuscript, we introduced `ROCCO`, a novel method for identifying open chromatin regions in ATAC-seq data that simultaneously leverages information from multiple samples to determine a consensus set of peaks. `ROCCO` uses spatial features of enrichment signal data by initially formulating the problem with a convex model that can be solved with provable efficiency and performance guarantees. Importantly, the model accounts for features common to the edges of accessible chromatin regions, which are often hard to determine based on independently determined sample peaks that can vary widely in their genomic locations. In addition to several attractive conceptual and theoretical features, `ROCCO` also exhibited improved detection performance based on ATAC-seq data from 56 lymphoblastoid samples evaluated against known transcription factor binding sites determined using ChIP-seq. `ROCCO` is especially suited for experimental designs that include samples from two or more distinct groups with one goal being to determine regions that are differentially accessible between these groups. A Jupyter notebook tutorial provides a step-by-step protocol for this with all necessary scripts provided on the GitHub repository.

For simplicity and to provide a conservative comparison with other methods, we ran `ROCCO` with the same genome-wide parameters for all chromosomes, including the budget which dictates the *maximum* proportion of the chromosome that should be considered accessible. However, chromatin accessibility varies across chromosomes, and `ROCCO`’s performance may be improved by exploiting properties specific to each chromosome. We found that optimizing the budget parameter for each chromosome for ℱ_*β*_-score at *β* = 1.0 did show an improvement. These optimized budget parameters roughly reflected the differences in gene density and read density across chromosomes, as expected. In the future, we will focus on developing an efficient, robust method to derive reasonable chromosome-specific budgets based on the input signal data.

`ROCCO`’s locus size parameter was set to *L* = 50 throughout experiments. While our results suggest this grants good performance and a resolution sufficient to identify both broad and concentrated regions of enrichment, it may prove beneficial to modify this parameter depending on the expected size of elements and desired granularity. We note, however, that decreasing *L* increases the number of loci, *n*, which may induce additional computational expense. By the same reasoning, computational burden can be reduced by increasing *L*, though some loss in the precision of predicted peaks may result.

Overall, `ROCCO` represents a scalable, effective, and mathematically sound method that is broadly applicable and addresses an important need in functional genomics analysis.

## 5 Methods

### 5.1 Computing Environment

Experiments were conducted using a stand-alone computer with an Intel Xeon CPU E5-2680 v3 @ 2.50GHz processor, 8 cores, and 64g RAM. The MOSEK solver [MOSEK, 2022], for which a free academic license can be readily obtained^3^, is used to solve (6). We run `rocco prep` to process BAM files and create enrichment signal tracks with *L* = 50.

### 5.2 Data Availability Statement

The ATAC-seq data from 56 lymphoblastoid samples used to conduct experiments was obtained from the ENCODE Project [Luo et al., 2019]. Specifically, we used alignments that had been determined according to the ENCODE ATAC-seq protocol. Note, we remove chromosome Y from consideration to ensure each sample contained data for the same set of chromosomes. A link to the metadata for this dataset is available in Supplementary Material 1.6.

## 1 Supplementary Material

### 1.1 Proof of Theorem I

For easy reference, the optimization problem (5) is restated here.

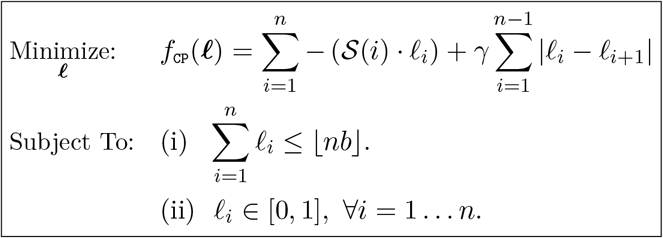

#### Theorem 1.

*(5) can be solved in polynomial time for a globally optimal solution*.

*Proof*. The crux of the proof is to show that (5) can be expressed as a linear program (LP): That is, a constrained optimization problem with real-valued decision variables in which the objective function and constraints are linear. These conditions maintain a convex feasible region of solutions with a unique (global) minimum value. Efficiency then follows from the use of interior-point algorithms with well-known polynomial time complexities for LPs [den Hertog, 1994].

A quick remedy for modelling the non-linearity of *f* is to introduce additional variables, *ℓ*_*j*_ :*∀i < n, j* = *n* + *i* to replace the absolute value functions in the objective

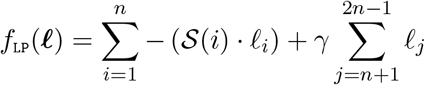

and add constraints such that:

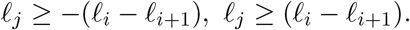

Since we are *minimizing f*, in any optimal solution, the above constraints imply

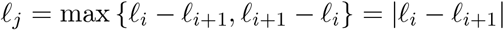

for all *ℓ*_*i*_. Thus, the truncated *n*-dimensional solution to the following LP:

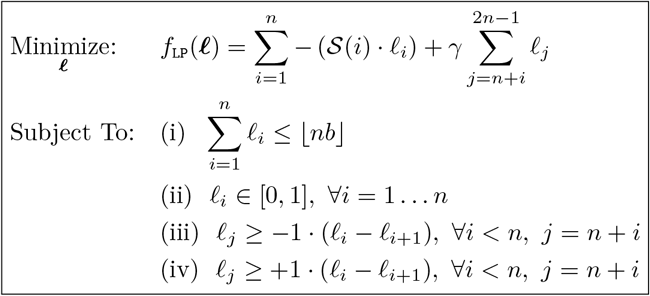

is an optimal solution for (5), which is what we needed to show. □

### 1.2 Proof of Theorem 2

The following two results are are useful in proving Theorem 2.

#### Lemma 1.

*Let* ***ℓ*** ^*rand*^ *be a solution generated with the procedure in Algorithm 1 and let* ***ℓ***^*LP*^ *be an optimal solution to 6. Then* ***ℓ*** ^*rand*^ *is integral and possesses the following properties*

1. *The objective value of* ***ℓ*** ^*rand*^ *is equal in expectation to the objective value of* ***ℓ*** ^*LP*^ :

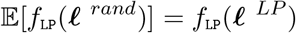
2. *The randomized solution satisfies the budget in expectation:*

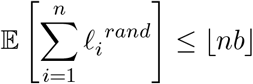

Both of the statements above follow as consequences of linearity of expectation.

Note, without loss of generality, we can treat the objective function *f* as non-negative in order to satisfy the condition for Markov’s inequality, used in proof of Theorem 2. Because *f* is bounded below, we can always add a constant to ensure min *f ≥*0 without affecting optimality of solutions since

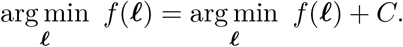

#### Theorem 2.

*Let* **L**_*N*_ *be a set of N ≥*1 *random solutions generated with Algorithm 1, and let c >* 1, *a >* 0 *be real numbers satisfying* 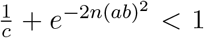 *for n loci and budget b∈*(0, 1). *Then with high probability*, **L**_*N*_ *contains at least one solution with both (a) an objective value no more than c·*OPT *and (b) no more than nb*(1 + *a*) *loci selected*.

*Proof*. To prove the result, we find a lower bound for the probability that these two criteria occur simultaneously for some solution in **L**_*N*_, and that this probability approaches 1 as *N → ∞* . Note, for concision, we let 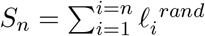.

Now, using the expected value for the objective given in Lemma 1, applying Markov’s inequality yields:

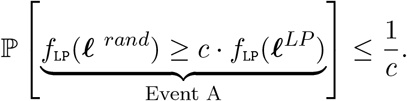

Likewise, recalling that each 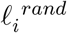 is generated as an independent random variable, we can apply Hoeffding’s inequality and Lemma 1 to obtain:

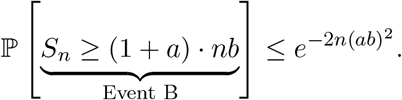

Combining these two (possibly dependent) events with the union bound, we get

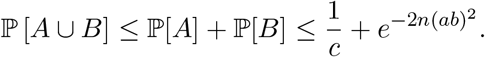

By construction, each solution in **L**_*N*_ is independent and identically distributed. Since the joint probability of *N* independent events *{E*_1_, *E*_2_, …, *E*_*N*_*}*, each satisfying ℙ [*E*_*i*_] *≤m*_*p*_, is no more than (*m*_*p*_)^*N*^, we can obtain an upper bound for the probability that at least one of the events, *A* or *B*, occurs in *every* solution in **L**_*N*_ :

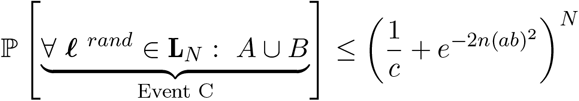

We are interested in the event that,

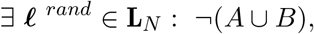

or, equivalently,

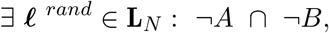

where at least one solution exists in **L**_*N*_ such that neither *A* nor *B* occur. This is the complement of Event *C* and occurs with probability at least

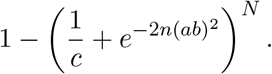

More explicitly, we have:

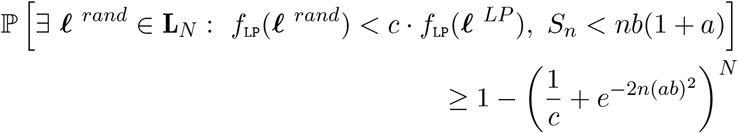

which approaches 1 for increasing *N* . □

### 1.3 Computational Efficiency

In this section, we discuss `ROCCO`’s computational efficiency from both theoretical and practical perspectives. Unless otherwise stated, `ROCCO`’s default parameters and the computing environment discussed in Section 5.1 are used.

The time complexity of `ROCCO` is equivalent to the time complexity required to solve a linear program (LP) with *n* variables and a *d*-bit data representation, since this step dominates the others asymptotically (See Algorithm 2). However, the 𝒪 (*n*^3^*d*) bound of standard interior-point solvers is of little insight for the vast majority of practical problems using current methodology and is only realized in pathological cases with completely dense matrices. A detailed survey of the improved capabilities of modern solvers and the multitude of algorithmic and hardware-based advances experienced in the past two decades is out of scope for this paper, but readers are referred to [Koch et al., 2022].

To gain a practical sense of `ROCCO`’s computational expense, an empirical analysis is warranted. To this end, we ran `ROCCO` genome-wide in 50 independent trials on the data set described in Section 5.2. The observed mean runtime and maximum resident set size (MRSS) for each chromosome were then plotted, with the 95%-confidence intervals^4^ thereof shaded to account for uncertainty arising from stochastic factors at the operating system and processor-level in Figure S1.

**Table S1:** Chromosome Sizes (bp) in hg38.

| Chromosome | Base pairs (Millions) |
| --- | --- |
| 1 | 248.96 |
| 2 | 242.19 |
| 3 | 198.30 |
| 4 | 190.21 |
| 5 | 181.54 |
| 6 | 170.81 |
| 7 | 159.35 |
| 8 | 145.14 |
| 9 | 138.39 |
| 10 | 133.80 |
| 11 | 135.09 |
| 12 | 133.28 |
| 13 | 114.36 |
| 14 | 107.04 |
| 15 | 101.99 |
| 16 | 90.34 |
| 17 | 81.28 |
| 18 | 78.08 |
| 19 | 59.13 |
| 20 | 63.03 |
| 21 | 48.13 |
| 22 | 50.96 |
| X | 155.27 |
| Y | 59.37 |

Chromosome size proved to be a fundamental contributor to computational expense. For example, `chr1`—the largest of human chromosomes—yielded the greatest average runtime and MRSS. In contrast, the smallest chromosome, `chr21`, required roughly one-tenth the time of `chr1`. See Table S1 for the number of base pairs in each human chromosome, scaled to millions.

**Figure S1:**
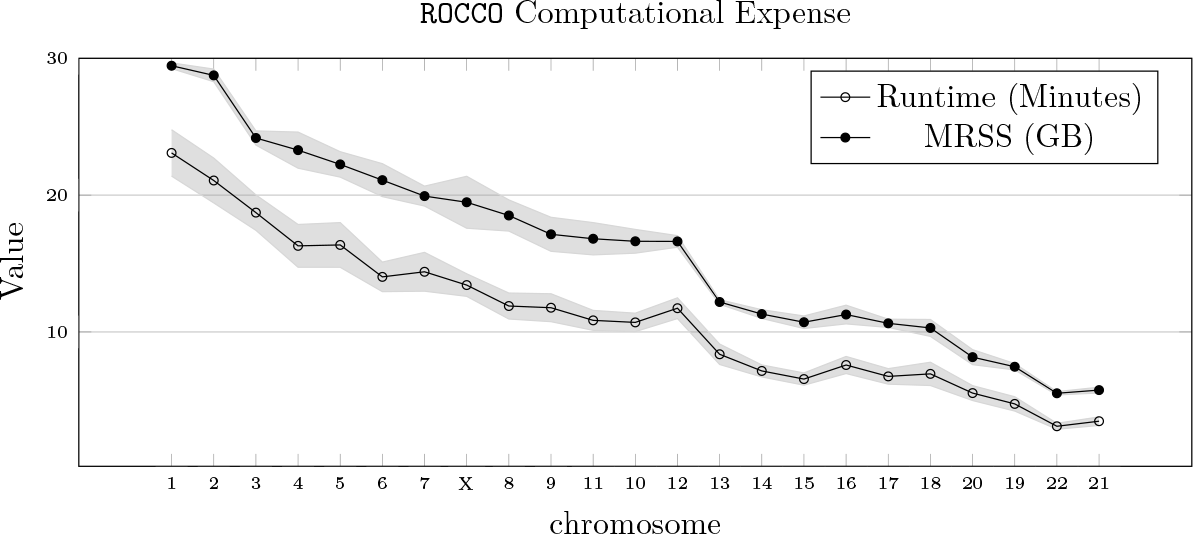
Average runtime and peak memory (MRSS) expended by `ROCCO` for each chromosome.

We note that `ROCCO`’s design allows for straightforward parallelization, since each chromosome can be processed independently. An option for simultaneous execution of `ROCCO` on multiple chromosomes is available in the software implementation via `rocco gwide --multi`. This feature can dramatically reduce the runtime of `ROCCO` if multiple cores are available. However, to accommodate users in limited computing environments, the default behavior of `rocco gwide` is to run the method sequentially on each chromosome, leading to an average genome-wide runtime of 4.3 hours in this experiment^5^.

### 1.4 ROCCO Parameters

`ROCCO` is designed to require minimal fine-tuning. Many of the parameters are included only for completeness and can be ignored for common use-cases. The default parameters provide generally strong performance as shown in Table S2, but we highlight several potentially consequential dynamics for users wishing to configure `ROCCO` optimally for a particular setting. In particular, the budget parameter may non-trivially affect behavior of the method.

#### 1.4.1 Constant Budgets

The budget, *b*, is a fundamental parameter of `ROCCO` as it determines an *upper-bound* on the proportion of loci that can be selected. A genome-wide, constant budget can yield strong detection performance as demonstrated in Tables 2, S2. The best constant budget depends on users’ preferences regarding recall/precision, as seen in Figure S2. Intuitively, for increasing *β*, that is, an increased emphasis on recall than precision, the best-scoring budget *b* is increasing.

**Figure S2:**
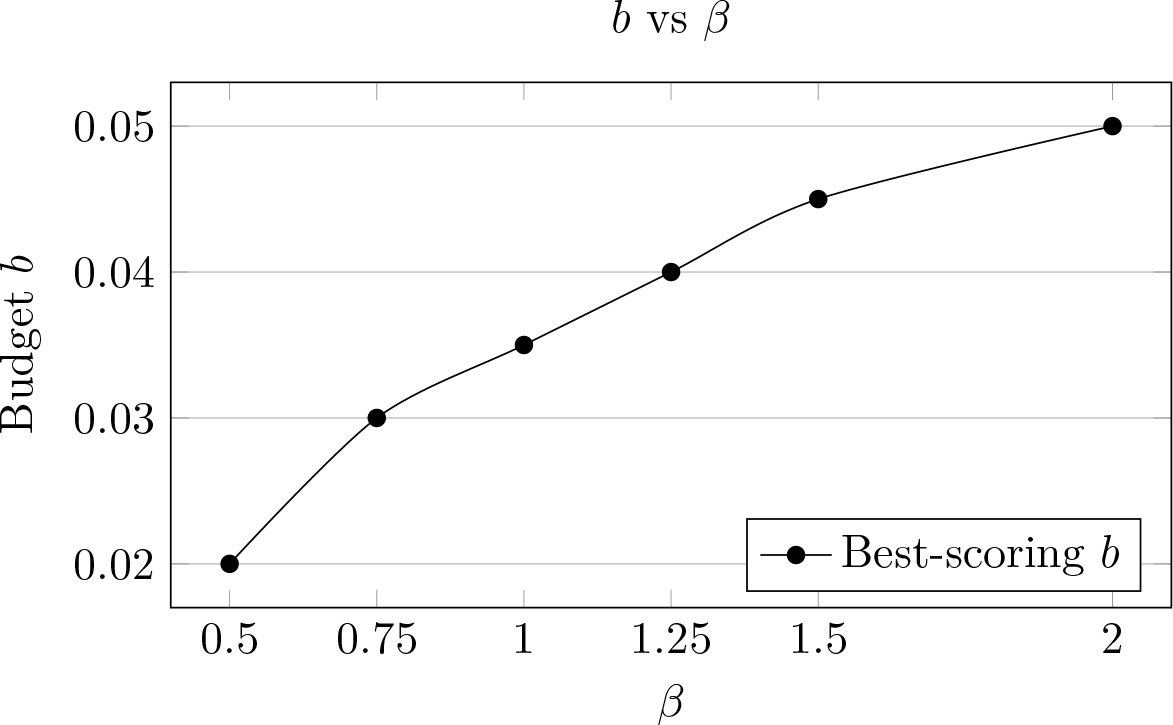
As greater importance is placed on recall than precision (*β* is increased), the ℱ_*β*_-tuned budget increases.

#### 1.4.2 Chromosome-Specific Budgets

A notion of dynamic, region-specific budgets is naturally appealing given chromatin’s variable dynamics throughout the genome. `ROCCO`’s software implementation allows users to specify budgets (and other parameters as well) at the chromosome level using `rocco gwide’s` `-p/--param_file parameter`. See hg38_params.csv for suggested budgets for each chromosome in the hg38 assembly. Using these chromosome-specific budgets to run `ROCCO` genome-wide on the ATAC-seq data in Section 5.2, ℱ_1_ = .592 was obtained as compared to ℱ_1_ = .579 with an invariant, genome-wide budget (Table 2). In the software implementation, `rocco budgets` can also be used to compute chromosome-specific budgets. In this procedure, estimates for the relative read densities of each chromosome are scaled such that they average to a user-specified value, e.g., *b* = .035. The ranking of chromosomes’ budgets according to read density is preserved under this operation, and the scaling step produces interpretable budget values as we have defined them. See [the API reference](https://nolan-h-hamilton.github.io/ROCCO/rocco/est_budgets.html) and [corresponding script](https://github.com/nolan-h-hamilton/ROCCO/blob/main/rocco/est_budgets.py) for more details.

#### 1.4.3 Locus Size, *L* and *γ*

At higher resolutions, it is natural to expect less variance between proximal elements, as the absolute distance between them is lesser by definition. Consequently, for smaller locus sizes *L*, users may wish to enforce a stronger dependence between adjacent loci by increasing *γ*. As a brief demonstration in Figure S3, all parameters except *γ, L* are fixed at their defaults, and we observe greater fragmentation in `ROCCO`’s solution with (*γ* = 1, *L* = 25) than the (*γ* = 2, *L* = 25), (*γ* = 1, *L* = 50), and (*γ* = 2, *L* = 50) cases.

**Figure S3:**
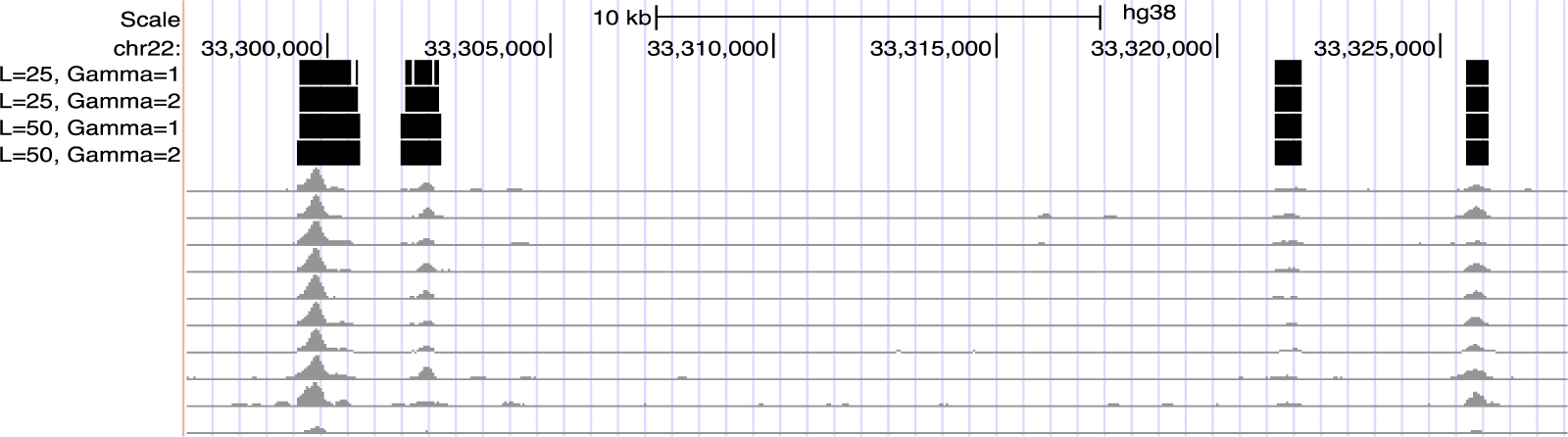
Relationship between γ and L. `ROCCO`’s annotations on a demonstrative 10kb subset of chromosome 22 are displayed. `ROCCO` is run four times on a set of ten ATAC-seq alignments with all parameters default/fixed except *γ, L*.

#### 1.4.4 Default Performance

**Table S2:** *ℱ*_*β*_-score obtained from each method using their default significance threshold.

| $\beta$ | ROCCO | Genrich | MACS2-Pooled | MACS2-Consensus |
| --- | --- | --- | --- | --- |
| 0.50 | 0.597 | 0.277 | 0.329 | **0.642** |
| 0.75 | **0.587** | 0.317 | 0.369 | 0.565 |
| 1.0 | **0.579** | 0.363 | 0.413 | 0.511 |
| 1.25 | **0.572** | 0.410 | 0.455 | 0.501 |
| 1.50 | **0.568** | 0.454 | 0.493 | 0.452 |
| 2.0 | 0.562 | 0.530 | **0.565** | 0.424 |

To measure “out-of-box” performance of `ROCCO`, we also compare each method using their *default* significance thresholds: *b* = .035, *q* = .05, *p* = .01 for `ROCCO`, MACS2, and Genrich, respectively. In Table S2, we see that `ROCCO` outperforms the benchmarks in four of six experiments. The two experiments in which `ROCCO` does not achieve best performance are for the minimum and maximum *β* values, where recall is weighed half and twice as much as precision, respectively. Users with a strong preference for either recall or precision are advised to tune accordingly.

### 1.5 Randomness of RR Procedure

Using the data set described in Section 5.2, `ROCCO` is run five times independently using the default *N* = 50 RR iterations. The pairwise Jaccard statistics between resulting peak sets are then computed in Table S3.

**Table S3:** Pairwise Jaccard indexes of peak sets obtained during five separate genome-wide runs of ROCCO with default parameters and *N* = 50 RR iterations.

| $\mathcal{J}$ | $\ell_1^*$ | $\ell_2^*$ | $\ell_3^*$ | $\ell_4^*$ | $\ell_5^*$ |
| --- | --- | --- | --- | --- | --- |
| $\ell_1^*$ | 1 | 0.997758 | 0.997745 | 0.997765 | 0.997743 |
| $\ell_2^*$ | . | 1 | 0.997734 | 0.997745 | 0.997714 |
| $\ell_3^*$ | . | . | 1 | 0.997721 | 0.997731 |
| $\ell_4^*$ | . | . | . | 1 | 0.997738 |
| $\ell_5^*$ | . | . | . | . | 1 |

#### 1.6 ATAC-seq Data

The 56 ATAC-seq samples used for experiments in this paper are available from the ENCODE Project. These BAM files result from ENCODE’s standard ATAC-seq protocol https://www.encodeproject.org/pipelines/ENCPL344QWT/.

Download metadata for ATAC-seq samples: https://www.encodeproject.org/metadata/?control_type%21=%2A&status=released&perturbed=false&assay_title=ATAC-seq&biosample_ontology.cell_slims=lymphoblast&files.file_type=bam&type=Experiment

We select the BAM files corresponding to rows with Output Type=‘alignments’.

### 1.7 Conservative IDR TF Chip-Seq Union Peaks

Available from: https://github.com/Boyle-Lab/F-Seq2-Paper-Supplementary/blob/d9c5357e615535240b09fac6567874bf6ATAC-seq%20benchmarking/data_process/download_preprocess_data.ipynb

We then used the textttliftOver UCSC binary tool genome.ucsc.edu/cgi-bin/hgLiftOver to convert to hg38.

### 1.8 Alternative Methods - Configurations

#### 1.8.1 MACS2-Based Methods

The following configuration of `MACS2` is used to call peaks in each sample for each significance threshold.

~~~
     macs2 callpeak -t {BAM file} -f BAMPE --nomodel --nolambda \
     --keep-dup all -q {significance threshold}
~~~

#### 1.8.2 Genrich

Genrich is called with the following configuration for each significance threshold

~~~
     ./Genrich -t {comma-separated list of name-sorted BAM files} -j \
     -p {significance threshold}
~~~

The `-j` option specifies ATAC-seq mode

### 1.9 Peak Width Distributions

Each method is tuned for the ℱ_1_-score as in Table 2 Row 3, and their respective peak-width distributions plotted in Figure S4. For each method, there is a consistent majority of peaks less than 500 bps with strictly decreasing frequency past this length.

**Figure S4:**
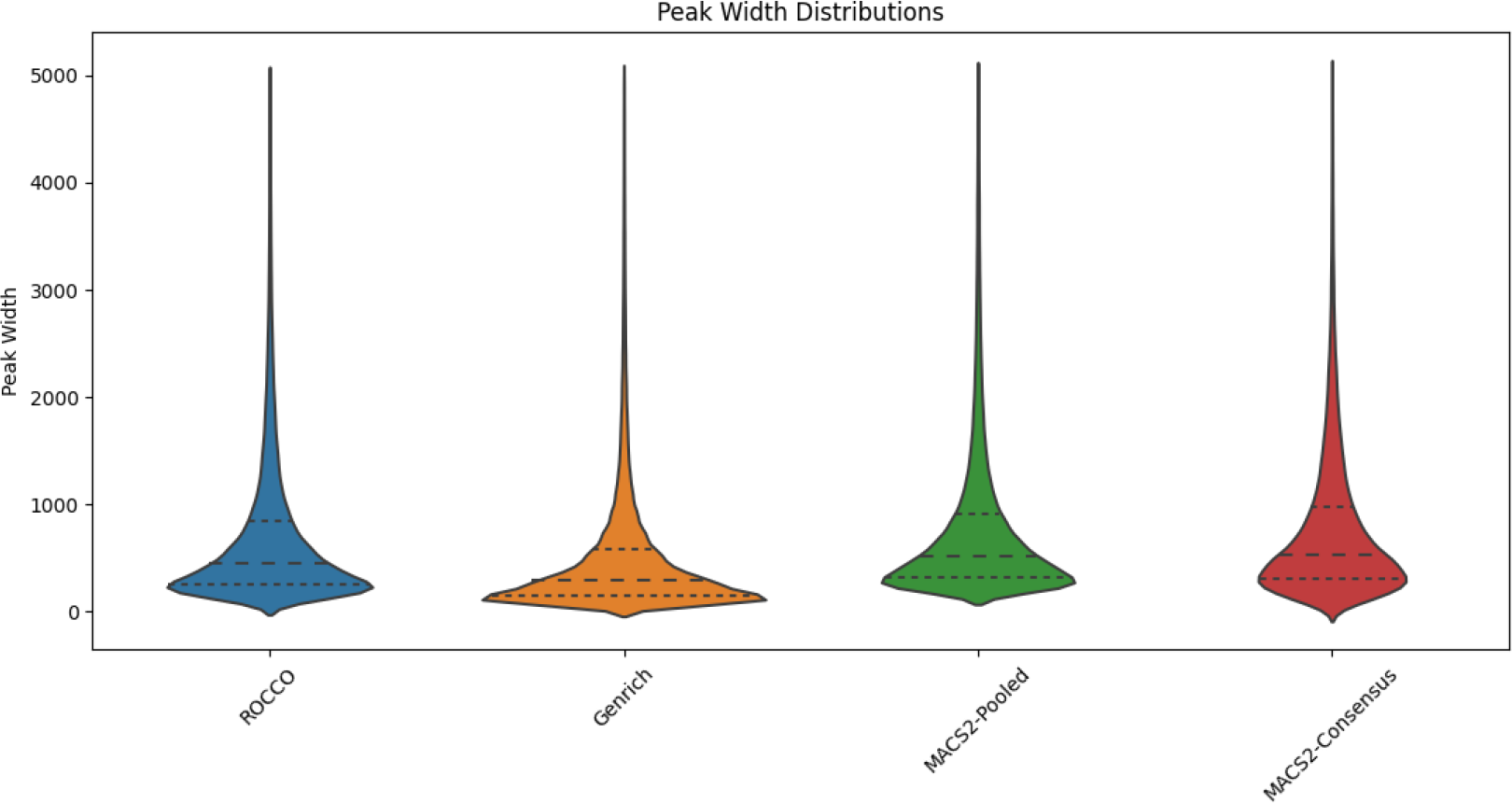
Violin plots of the peak width distributions for each method.

## Footnotes

1 Available at https://github.com/jsh58/Genrich

2 Supplementary Material 1.2

3 `ROCCO` can call any open-source LP solver offered within the CVXPY [Diamond and Boyd, 2016] platform, but runtimes may vary. ECOS [Domahidi et al., 2013] is a viable option installed with CVXPY by default.

4 The confidence intervals were computed according to a *t*-distribution with *v* = 49.

5 In a personal computing environment, `rocco gwide` successfully returned genome-wide results on a MacBook ([Model A2485](https://support.apple.com/kb/SP858?locale=en_US)) in under five hours.

